# Analysis of the SARS-CoV-2 spike protein glycan shield: implications for immune recognition

**DOI:** 10.1101/2020.04.07.030445

**Authors:** Oliver C. Grant, David Montgomery, Keigo Ito, Robert J. Woods

## Abstract

Here we have generated 3D structures of glycoforms of the spike (S) glycoprotein from SARS-CoV-2, based on reported 3D structures and glycomics data for the protein produced in HEK293 cells. We also analyze structures for glycoforms representing those present in the nascent glycoproteins (prior to enzymatic modifications in the Golgi), as well as those that are commonly observed on antigens present in other viruses.

These models were subjected to molecular dynamics (MD) simulation to determine the extent to which glycan microheterogeneity impacts the antigenicity of the S glycoprotein. Lastly, we have identified peptides in the S glycoprotein that are likely to be presented in human leukocyte antigen (HLA) complexes, and discuss the role of S protein glycosylation in potentially modulating the adaptive immune response to the SARS-CoV-2 virus or to a related vaccine.

The 3D structures show that the protein surface is extensively shielded from antibody recognition by glycans, with the exception of the ACE2 receptor binding domain, and also that the degree of shielding is largely insensitive to the specific glycoform. Despite the relatively modest contribution of the glycans to the total molecular weight (17% for the HEK293 glycoform) the level of surface shielding is disproportionately high at 42%.

## Introduction

The present COVID-19 pandemic has led to over a million confirmed infections globally with a fatality rate of approximately 5 percent (1) since the first reports of a severe acute respiratory syndrome (SARS) infection by a novel coronavirus (SARS-CoV-2) at the end of 2019. As of April 2020, there is still no vaccine or approved therapeutic to treat this disease. Here we examine the structure of the SARS-CoV-2 envelope spike (S) protein that mediates host cell infection, with a specific focus on the extent to which glycosylation masks this virus antigen from the host immune response.

Viral envelope proteins are often modified by the attachment of complex glycans that can account for up to half of the molecular weight of these glycoproteins, as in HIV gp120 (2). The glycosylation of these surface antigens helps the pathogen evade recognition by the host immune system by cloaking the protein surface from detection by antibodies, and can influence the ability of the host to raise an effective adaptive immune response (3, 4) or even be exploited by the virus to enhance infectivity (5). Additionally, because the virus hijacks the host cellular machinery for replication and subsequent glycosylation, the viral glycan shield may be composed of familiar host glycans; thereby suppressing an anti-carbohydrate immune response (6).

Fortunately, the innate immune system has evolved a range of strategies for responding to glycosylated pathogens (7), but antigen glycosylation nevertheless complicates the development of vaccines (8). Over time, the protein sequences in viral antigens undergo mutations (antigenic drift), which can alter the species specificity of the virus (9), modulate its infectivity (10), and alter the antigenicity of the surface proteins (11). These mutations can also impact the degree to which the protein is glycosylated by creating new or removing existing locations of the glycans (glycosites) on the antigens (12, 13). Varying surface antigen glycosylation is thus a mechanism by which new virus strains can evade the host immune response (12), and attenuate the efficacy of existing vaccines (8).

Very recently, a cryo-EM structure of the SARS-CoV-2 S glycoprotein has been reported (14), which led to conclusion that, like the related protein from the 2002 - 2003 SARS pandemic (SARS-CoV-1) (15), the CoV-2 S protein is also extensively glycosylated (14). Furthermore, an analysis of the glycan structures present at each glycosite in the S protein produced recombinantly in human embryonic kidney (HEK) 293 cells has also been recently reported (16).

Here we have generated 3D structures of several glycoforms of the SARS-CoV-2 S glycoprotein, in which the glycans represent those present in the S protein produced in HEK293 cells (16), as well as those corresponding to the nascent glycoprotein (prior to enzymatic modifications in the Golgi apparatus), and those that are commonly observed on antigens present in other viruses (17–19). We have subjected these models to long molecular dynamics (MD) simulations and compared the extent to which glycan microheterogeneity impacts epitope exposure. Additionally, we have identified peptides in the S protein that are likely to be presented in human leukocyte antigen (HLA) complexes, and discuss the role of S protein glycosylation in modulating the adaptive immune response to the SARS-CoV-2 virus or to a related vaccine.

The impact of glycosylation on the ability of antibodies to bind to a pathogenic glycoprotein may be estimated by quantifying the fraction of the surface area of the protein antigen that is physically shielded by glycans from antibody recognition. However, glycans display large internal motions that prevents their accurate description by any single 3D shape, in contrast to proteins (20, 21). Fortunately, MD simulations can play a key role by accurately predicting the 3D shapes and motions of glycans, as confirmed by comparison to solution NMR data (22–24), and such simulations have been widely applied to glycoproteins (17, 25–28). Here we have performed MD simulations with the GLYCAM06/AMBER force field, which was developed for simulations of carbohydrates, carbohydrate-protein complexes and glycoproteins (29–31), and use the data to assess the impact of glycosylation on the immunogenic and antigenic properties of the S glycoprotein.

## Results

### Model glycoforms

It is well established that there is a strong dependence of both the composition and relative glycan abundance (glycan microheterogeneity) on the cell type used for glycoprotein production. And there is a large body of data relating to the influence of host cell line on viral envelop protein glycosylation. For example, a glycomics analysis of influenza A virus produced in five different cell lines, all of relevance to vaccine production, led to the observation of profound differences in the compositions of the glycans at a given site; with structures varying from paucimannose (Sf9 cells) to core-fucosylated hybrid with bisecting N-acetylglucosamine (Egg) to sialylated biantennary glycans (HEK293) (19). For these reasons, we have modeled the S glycoprotein with reported site-specific glycosylation (16), as well as hypothetical homogeneously glycosylated glycoforms of the high mannose (M9), paucimannose (M3), biantennary complex (Complex) and core-fucosylated biantennary complex (Complex Core F) types. Comparisons among the glycoforms permits an assessment of the impact of cell-based differential glycan processing on S protein antigenicity.

### Assessment of the impact of glycosylation on antigenicity

We subjected the five glycoforms of the CoV-2 S glycoprotein to MD simulation and interpreted the results in terms of the impact of glycan structure on the theoretical S glycoprotein antigenic surface area (Figure 1, Supplementary Figure S1, Table 1). A series of 3D structure snapshots of the simulation were taken at 1 ns intervals and analysed in terms of their ability to interact with a spherical probe based on the average size of hypervariable loops present in an antibody complementarity determining region (CDR) (Figure 3). The percentage of simulation time each residue was exposed to the antibody accessible surface area (AbASA) probe was calculated and plotted onto both the 3D structure (Figure 1) and sequence (Figure 2). The average AbASA over the course of the simulations was also calculated for each glycoform and compared to non-glycosylated protein (Table 1).

**Table 1.**
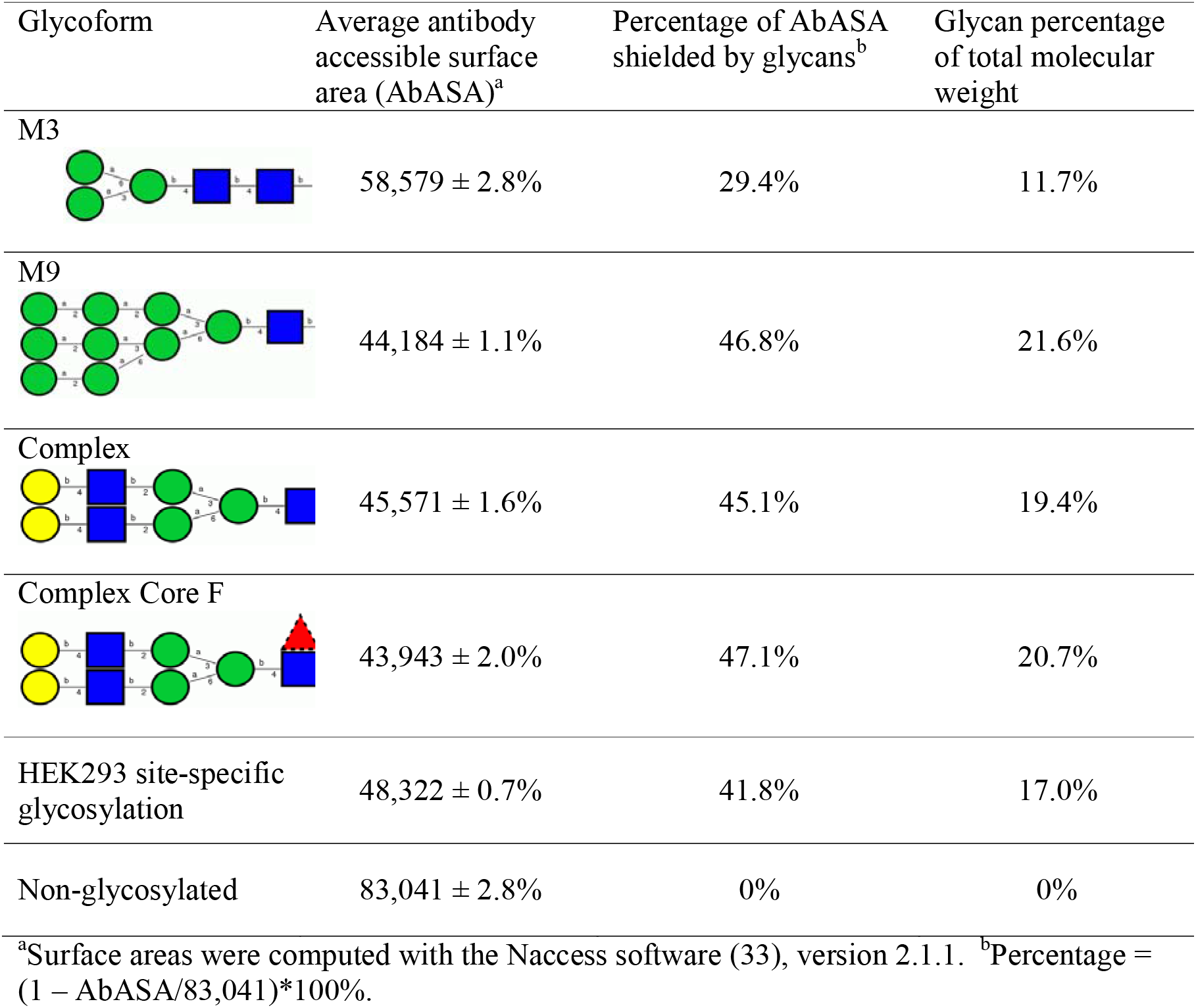
SARS-CoV-2 S glycoprotein antigenic surface areas (Å^2^) as a function of glycoform.

| Glycoform | Average antibody accessible surface area (AbASA)<sup>a</sup> | Percentage of AbASA shielded by glycans<sup>b</sup> | Glycan percentage of total molecular weight |
| --- | --- | --- | --- |
| M3 | 58,579 ± 2.8% | 29.4% | 11.7% |
| M9 | 44,184 ± 1.1% | 46.8% | 21.6% |
| Complex | 45,571 ± 1.6% | 45.1% | 19.4% |
| Complex Core F | 43,943 ± 2.0% | 47.1% | 20.7% |
| HEK293 site-specific glycosylation | 48,322 ± 0.7% | 41.8% | 17.0% |
| Non-glycosylated | 83,041 ± 2.8% | 0% | 0% |

<sup>a</sup>Surface areas were computed with the Naccess software (33), version 2.1.1. <sup>b</sup>Percentage = $(1 - \text{AbASA}/83,041) \times 100\%$ .

**Figure 1.**
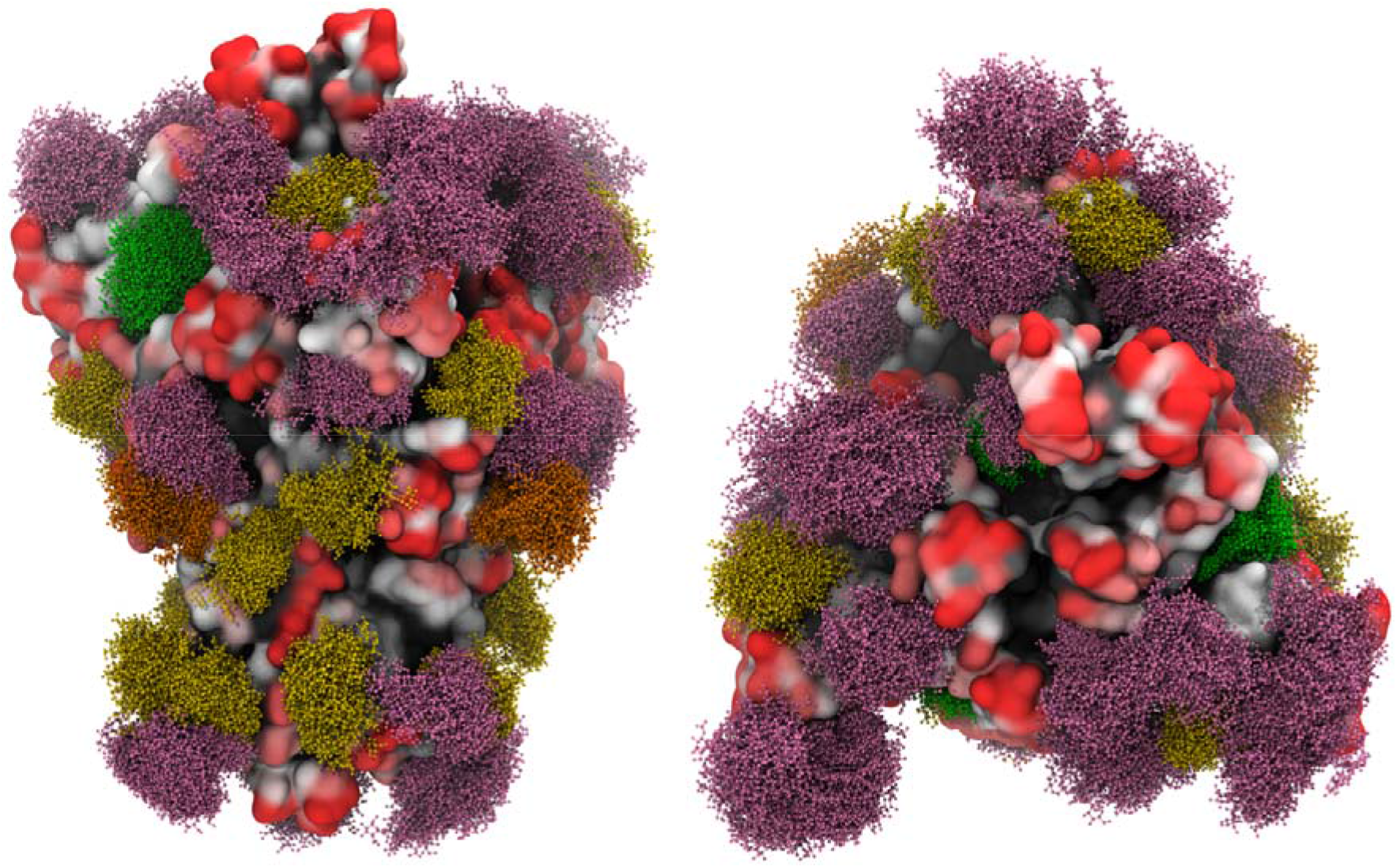
Overlay of snapshots from MD simulation of the S glycoprotein with site-specific glycosylation. The glycans are shown in ball-and-stick representation: M9 (green), M5 (dark yellow), hybrid (orange), complex (pink) (See Supplementary Table S1 for details). The protein surface is colored according to antibody accessibility from black to red (least to most accessible). Images generated using Visual Molecular Dynamics (VMD) (32) version 1.9.3.

**Figure 2.**
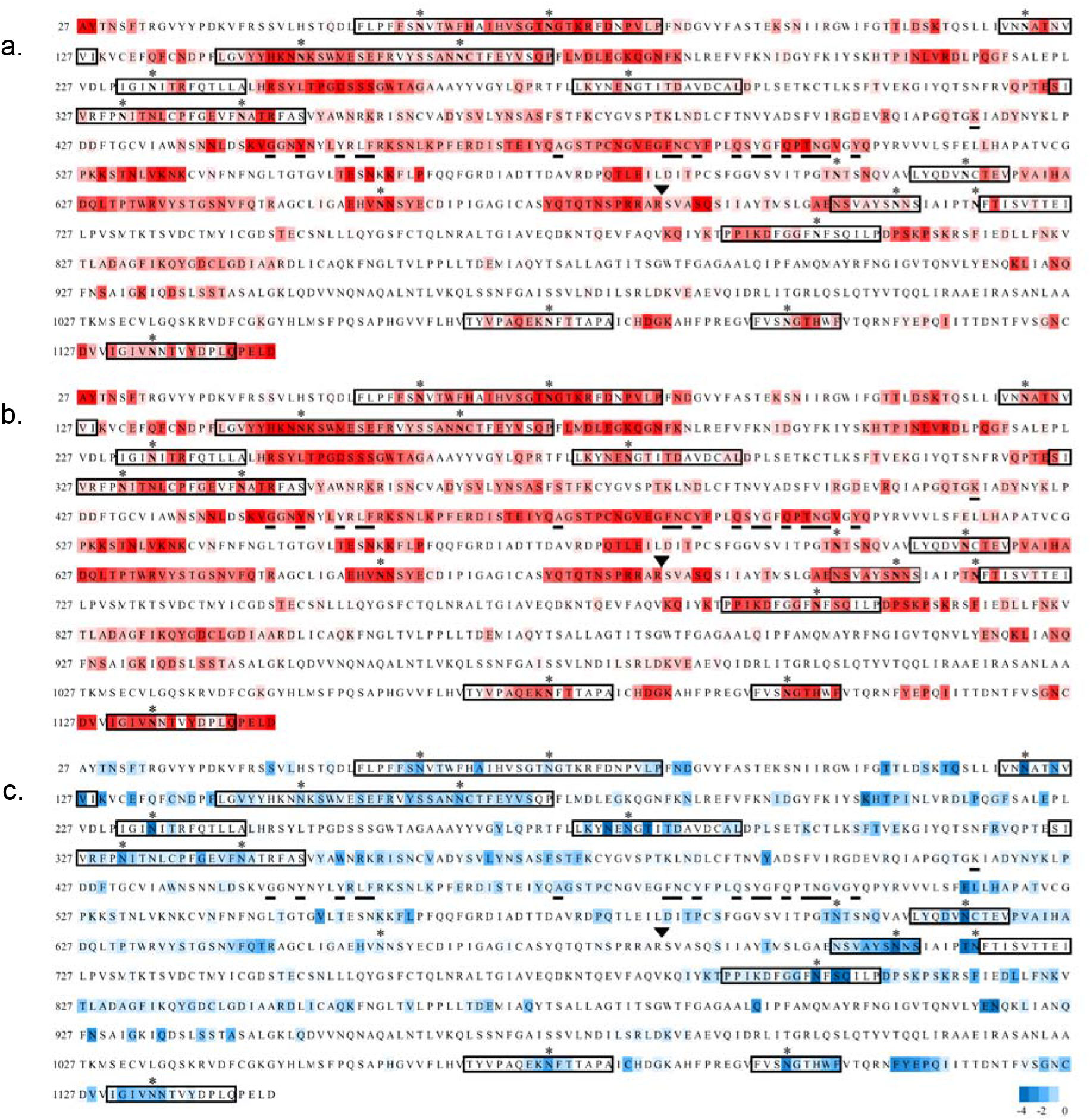
Sequence of the S protein (NCBI: YP_009724390.1) used to generate the 3D model of the glycoprotein. Residues 1-26 and 1147-1273 were not included in the 3D structure due to a lack of relevant template structures. Sequences within a rectangle were predicted to consist of one or more HLA antigens using the RankPep server (imed.med.ucm.es/Tools/rankpep (36, 37)). Glycosites are indicated with asterisks, residues reported to interact with the ACE2 receptor (42) are underlined, and the protease cleavage site is indicated with a triangle above the RS junction. **a.** The sequence is colored according to antibody accessibility computed for the site-specific glycoform from white to red (least to most accessible). **b**. Antibody accessibility computed for the non-glycosylated protein. **c**. The difference in accessibilities between the site-specific and non-glycosylated glycoforms is plotted as the fold change in epitope accessibility during the simulation, from −4 (blue) to 0 (white), where blue indicates glycosylation-dependent surface shielding.

**Figure 3.**
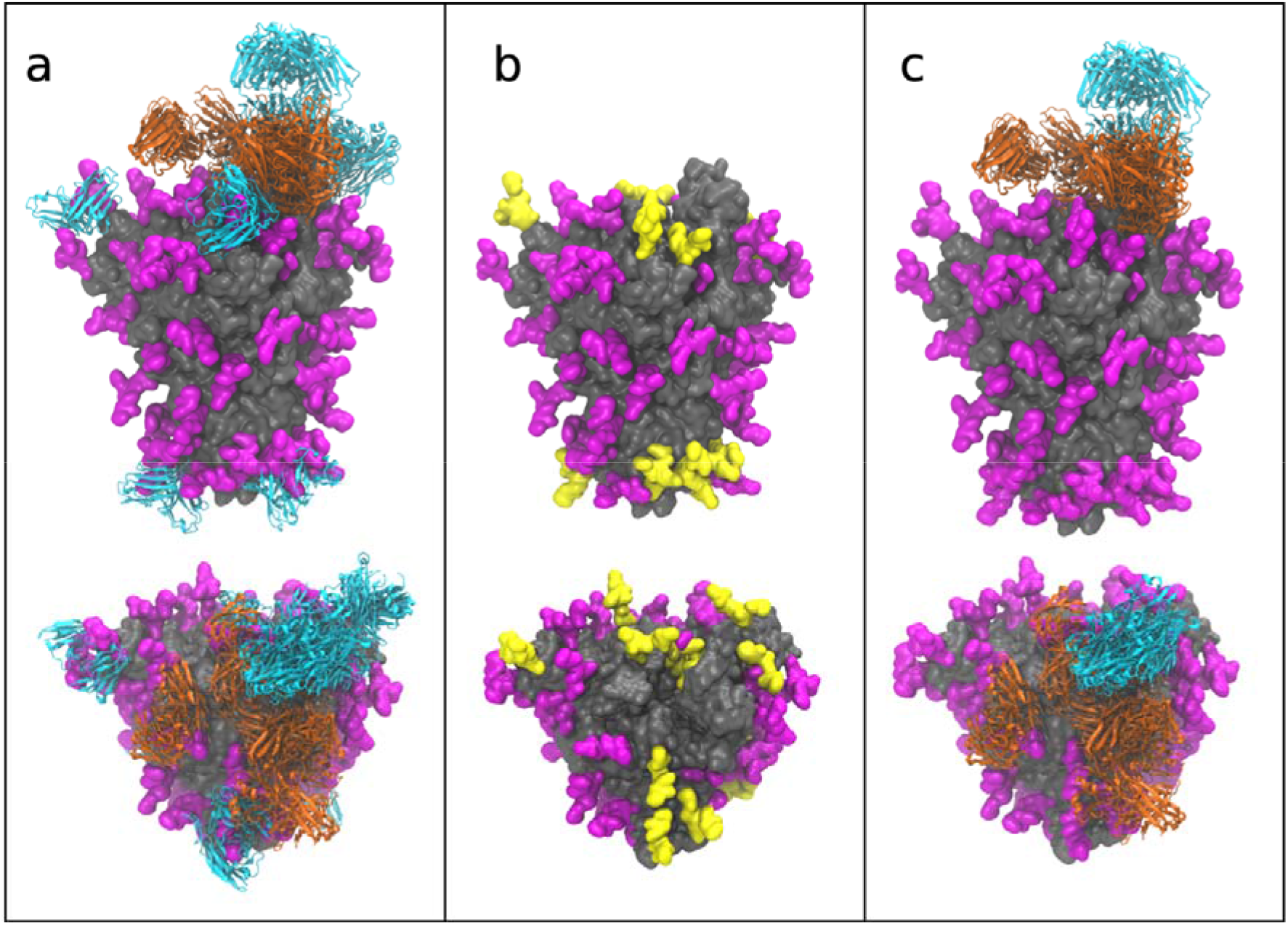
Glycan shielding of the SARS-CoV-2 S protein from known antibodies for the SARS-CoV-1 and MERS S proteins. **a.** Side view (upper panels) and top view (lower panels) of the SARS-CoV-2 S protein (grey surface) with homogeneous complex glycosylation (magenta) showing aligned antibody fragments (ribbons) from co-complexes with the S glycoproteins from SARS-CoV-1 (orange) and MERS (cyan) (44–56). **b.** Glycans present on SARS-CoV-2 S glycoprotein that are incompatible with known antibody positions due to steric overlap are shown in yellow. **c.** Potential antibody poses after elimination of epitopes blocked by S protein glycosylation. Images generated using VMD (32) version 1.9.3.

The data indicate that uniform glycosylation with the smallest of the glycans (paucimannose, M3), which is a sub-structure within all N-linked glycans, provided the least shielding of the S protein (29% coverage), leaving 71% of the surface exposed to an antibody probe relative to the same protein with no glycosylation. In contrast, the largest high mannose N-linked glycans (M9), which corresponds to the nascent glycoform that would exist prior to processing through the Golgi apparatus, led to the highest level of surface shielding (47%). The level of cloaking offered by the two types of complex glycans are not significantly different from that of M9 at 45-47% surface shielding. Glycosylation based on the S glycoprotein produced recombinantly in HEK293 resulted 42% of the surface being shielded from antibody recognition; a value that is comparable to that for the models based on uniform M9 or Complex glycans. There is a strong correlation (R^2^ = 0.98) between the total molecular weight of the glycans and the net shielding of the protein surface, however it is notable that the glycans disproportionately block the accessibility of the surface to the antibody probe. This result is indicative of the high density of the glycans on the S glycoprotein surface.

The results from the CDR accessibility analysis are consistent with the conclusion that antigenicity of the S protein is largely insensitive to glycan microheterogeneity, with the exception of the glycoform composed solely of M3 glycans. Nevertheless, differences in glycosylation may impact other structural features, such as local interactions between the glycan and the protein surface, or local structural fluctuations in either the protein or glycan conformations that are only partially captured by the exposed surface area analysis.

A visual examination of the glycoform 3D structures (Figure 1 and Supplementary Figure S1) indicates that the most exposed epitopes comprise the ACE2 receptor site, specifically the apex region of the S1 domain when that domain is in the open conformation. The large extent of RBD exposure is quantified in Figure 2c. Moreover, the extensive motion displayed by each glycan illustrates that no single static model could fully capture the extent of glycan shielding. It can also be observed that a ring of antigenic sites appears to encircle the S1 domain, independent of glycoform. Unlike the extremely high level of glycan shielding in gp120 that challenges HIV vaccine development (34, 35), the level of shielding by glycans in the S protein is more moderate, with approximately 42% of the surface potentially inaccessible to antibodies.

### Adaptive immune response to SARS-CoV-2

Beyond a role in shielding the underlying protein from recognition by antibodies, the glycans on pathogenic proteins may also attenuate the ability of the host immune system to raise antibodies against any epitopes that include the glycan. In a T-cell dependent adaptive immune response, peptides from the pathogen are presented on antigen presenting cells by major histocompatibility complex II molecules, known as human leukocyte antigen (HLA) complexes. HLA complexes have preferred peptide motifs, and based on a knowledge of these preferences it is possible to predict which peptides in a protein are likely to be HLA antigens (36, 37). However, when that peptide contains a glycosylation site, the ability of the peptide to be presented by an HLA may be compromised, if for example the peptide cannot bind to the HLA molecule due to the steric presence of the glycan. However, glycopeptides may be presented in HLA complexes if the glycan is small enough (38) or if it is found on the end of the peptide antigen where it doesn’t interfere with HLA binding (39). The glycan-mediated shielding of predicted HLA antigens (Supplementary Table S2) derived from the S protein are shown in Figure 4 and Supplementary Figures S2 and S3 for all HLA peptide sequences that also contain a glycosite.

**Figure 4.**
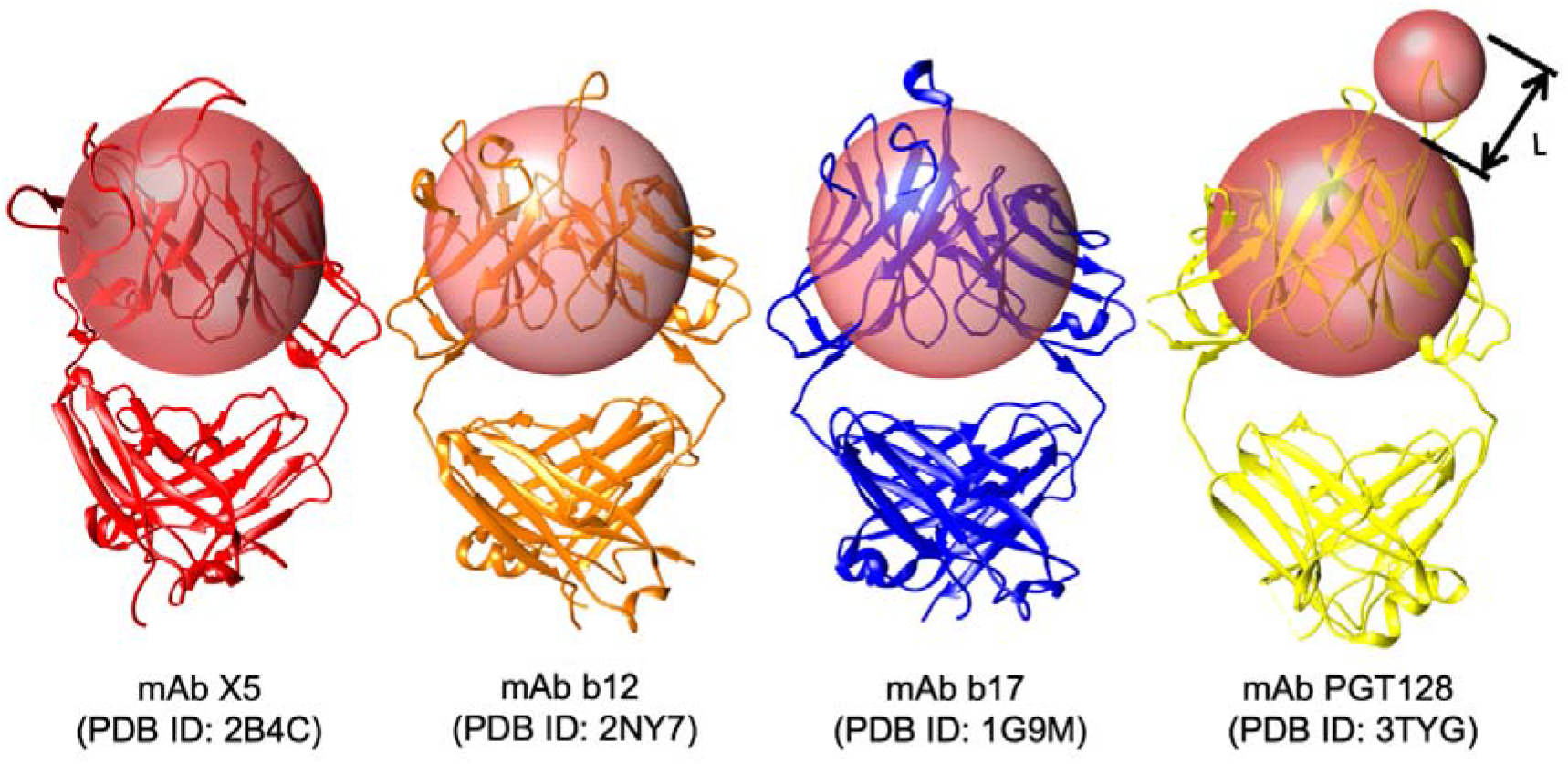
Antibody accessible surface area estimation using a pair of spherical probes. To estimate the AbASA, a spherical probe was derived (radius 7.2 Å, smaller sphere) that approximates the average size of the hypervariable loops from four anti-gp120 antibodies, in which the epitopes were either protein surface residues (PDB IDs: 2B4C (73), 2NY7 (74), 1G9M (75)) or both carbohydrate and protein residues: (3TYG (76)). This probe size may be compared to values of 5 and 10 Å employed previously to estimate antigenic surface area (77). Changes in the solvent accessible surface area (SASA) showed no shielding by glycans and thus a simple SASA model was not useful for this analysis (77). Additionally, to account for the presence of the beta-sheet framework in the antibody variable fragment (Fv), we introduced a second larger probe (18.6 Å) sufficient to approximately enclose that domain. The antigenic surface area is then defined as sum of the surface areas of any protein residues that make contact with the CDR probe, provided that the CDR probe is proximal to the Fv probe. This latter requirement is governed by “L”, which requires that the distance between the CDR-antigen contact site and the Fv probe surface be less than the length (10.4 Å) of the longest CDR loop in mAb PGT128. PGT128 was chosen for this reference as it contains a particularly long CDR loop that penetrates the glycan shield of gp120. Images generated with UCSF Chimera (78).

As expected, glycosylation consistently decreased the surface exposure of the residues proximal to the glycosites (Figure 2c), but also led to non-sequential changes in exposure, as a result of the 3D topology in the vicinity of each glycosite. Of the 18 glycosites in the 3D structure, 16 are predicted to be present in HLA peptides. Although the glycans may occur throughout the HLA sequences (Supplementary Table S2), in 12 of these sequences the glycans are predicted to be present at the terminus of at least one putative HLA antigen. This observation suggests that these 12 glycosites may not interfere with antigen presentation in an HLA complex. This property is essential for the potential generation of antibodies against the underlying epitopes, but moreover, may lead to antibodies that target these carbohydrates on the S glycoprotein (38). Anti-carbohydrate antibodies have been shown to be neutralizing in other viruses, such as HIV (40), and therefore glycosylated peptides can offer an alternative to more traditional peptide epitopes. Targeting glycans as epitopes is most effective when the glycans or their clusters are significantly different from self, and thus are not immunologically tolerated (6). Although viruses exploit the host glycosylation machinery in their biosynthesis, differences from host glycan distributions can occur when for example the virus cloaks itself so densely in glycosites that the glycans are not accessible to glycan processing enzymes, due to steric crowding, and remain in their high mannose form (17). Examples of this are seen in the high-mannose clusters in some strains of influenza (17) and in HIV (41).

From the perspective of vaccine development (43), targeting glycans as epitopes would be expected to benefit from matching the glycan microheterogeneity in the vaccine to that in the circulating virus, which requires additional consideration of the choice of cell type for vaccine production.

### Comparison with epitopes in related coronavirus S glycoproteins

To illustrate the impact of glycosylation on epitope exposure, we aligned the 3D structure of the spike proteins from SARS-CoV-2 with those from co-crystal structures of SARS-CoV-1 and MERS that contained bound antibody fragments. The S glycoproteins of SARS-CoV-1 and CoV-2 share a high degree of structural similarity, with an average root-mean-squared difference (RMSD) in the Cα positions of only 3.8 Å (14). The MERS S glycoprotein also shares a similar trimeric structure with CoV-1 and CoV-2. From this alignment, the extent to which epitopes in the CoV-2 S glycoprotein might be inaccessible to known antibodies on the basis of structural differences in the glycoproteins or due to shielding by glycans on the CoV-2 S glycoprotein surface was inferred (Figure 3). Approximately 50% of the corresponding epitopes in the CoV-2 S glycoprotein are blocked by glycans from antibody binding, and only areas of the protein surface at the apex of the S1 domain appear to be accessible to known antibodies. Although this static epitope analysis doesn’t take into account the plasticity of the glycans or the protein it does confirm that the RBD in the CoV-2 S glycoprotein should be accessible for antibody recognition, consistent with the analysis of the MD simulation data.

## Discussion

The present study indicates that glycans shield approximately 40% of the underlying protein surface of the S glycoprotein from antibody recognition, and that this value is relatively insensitive to glycan type. This suggests that although glycan microheterogeneity varies according to host cell type, the efficacy of antisera should not be impacted by such differences. In contrast, by analogy with influenza hemagglutinin (57, 58), variations in glycosite location arising from antigenic drift can be expected to have a profound effect on S protein antigenicity and potentially vaccine efficacy. Fortunately, the most accessible and largest epitope in the S protein consists of the ACE2 binding domain, where the virus cannot exploit glycan shielding or mutational changes to evade host immune response without potentially attenuating viral fitness. The requirement that the virus maintain the integrity of the ACE2 RBD suggests that a vaccine that includes this epitope may maintain efficacy despite antigenic drift, as long as the virus continues to target the same host receptor.

While overall shielding of the underlying protein surface does not appear to be highly sensitive to glycan microheterogeneity, it would likely impact the innate immune response by altering the ability of collectins and other lectins of the immune system to effectively bind to the S protein and neutralize the virus (17), and may impact the adaptive immune response by altering the number of viable HLA antigens. Given that in humans, glycan microheterogeneity varies between individuals, and depends on many factors, including age (59), underlying disease (60, 61) and ethnicity (62), access to 3D models of the S glycoprotein may aid in defining the molecular basis for the differential susceptibilities among individuals to COVID-19.

Lastly, the observation that homogeneously glycosylated glycoforms are predicted to display approximately the same shielding properties as those computed for the more relevant site-specific glycoform suggests that such models can be usefully applied in advance of the report of experimental glycomics data. This final conclusion is significant as it enables the effects of glycosite alterations to be estimated in anticipation of seasonal antigenic drift.

## Methods

### SARS-CoV2 spike (S) protein structure

A 3D structure of the prefusion form of the S protein (RefSeq: YP_009724390.1, UniProt: P0DTC2 SPIKE_SARS2), based on a Cryo-EM structure (PDB code 6VSB) (14), was obtained from the SWISS-MODEL server (swissmodel.expasy.org). The model has 95% coverage (residues 27 to 1146) of the S protein.

### S protein glycoform generation

Five unique 3D models for the glycosylated glycoprotein were generated using the glycoprotein builder available at GLYCAM-Web (www.glycam.org) together with an in-house program that adjusts the asparagine side chain torsion angles and glycosidic linkages within known low-energy ranges (63) to relieve any atomic overlaps with the core protein, as described previously (57, 64). The site specific glycans used to model a glycoform representative of the data obtained from the S glycoprotein expressed in HEK293 cells (16), are presented in Supplementary Table S1.

### Energy minimization and Molecular dynamics (MD) simulations

Each glycosylated structure was placed in a periodic box of approximately 130,000 TIP3P water molecules (65) with a 10 Å buffer between the glycoprotein and the box edge. Energy minimization of all atoms was performed for 20,000 steps (10,000 steepest decent, followed by 10,000 conjugant gradient) under constant pressure (1 atm) and temperature (300 K) conditions. All MD simulations were performed under nPT conditions with the CUDA implementation of the PMEMD (66, 67) simulation code, as present in the Amber14 software suite (68). The GLYCAM06j force field (69) and Amber14SB force field (70) were employed for the carbohydrate and protein moieties, respectively. A Berendsen barostat with a time constant of 1 ps was employed for pressure regulation, while a Langevin thermostat with a collision frequency of 2 ps^−1^ was employed for temperature regulation. A nonbonded interaction cut-off of 8 Å was employed. Long-range electrostatics were treated with the particle-mesh Ewald (PME) method (71). Covalent bonds involving hydrogen were constrained with the SHAKE algorithm, allowing an integration time step of 2 fs (72) to be employed. The energy minimized coordinates were equilibrated at 300K over 400 ps with restraints on the solute heavy atoms. Each system was then equilibrated with restraints on the Cα atoms of the protein for 1ns, prior to initiating 3 independent production MD simulations with random starting seeds for a total time of 0.75 μs, with no restraints applied.

### Antigenic surface analysis

A series of 3D structure snapshots of the simulation were taken at 1 ns intervals and analysed in terms of their ability to interact with a spherical probe based on the average size of hypervariable loops present in an antibody complementarity determining region (CDR) (Figure 3). The percentage of simulation time each residue was exposed to the AbASA probe was calculated and plotted onto both the 3D structure (Figure 1) and sequence (Figure 2). The average AbASA over the course of the simulations was also calculated for each glycoform and compared to nude protein (Table 2).

## Supporting Information

Coordinates in pdb format for each glycoform of the S glycoprotein are available for download from GLYCAM-Web (www.glycam.org).

## Acknowledgments

R.J.W. thanks the National Institutes of Health (U01 CA207824 and P41 GM103390) for financial support.

## Author Contributions

O.C.G. and R.J.W. designed the research; O.C.G., D. M., K. I., and R.J.W. performed the research; R.J.W. wrote the paper.

