## Supplementary Information for "Analysis of the SARS-CoV-2 spike protein glycan shield: implications for immune recognition"

*Mailing address: 315 Riverbend Road, Athens, GA 30602.

**Supplementary Figure S1.** Overlay of snapshots from MD simulation of the S glycoprotein with homogeneous glycosylation. The glycans are shown in ball-and-stick representation: M9 (green), M3 (cyan), complex and complex core F (pink) (See Table S1 for details). The protein surface is colored according to antibody accessibility from black to red (least to most accessible). Images generated using Visual Molecular Dynamics (VMD) ^62^ version 1.9.3.

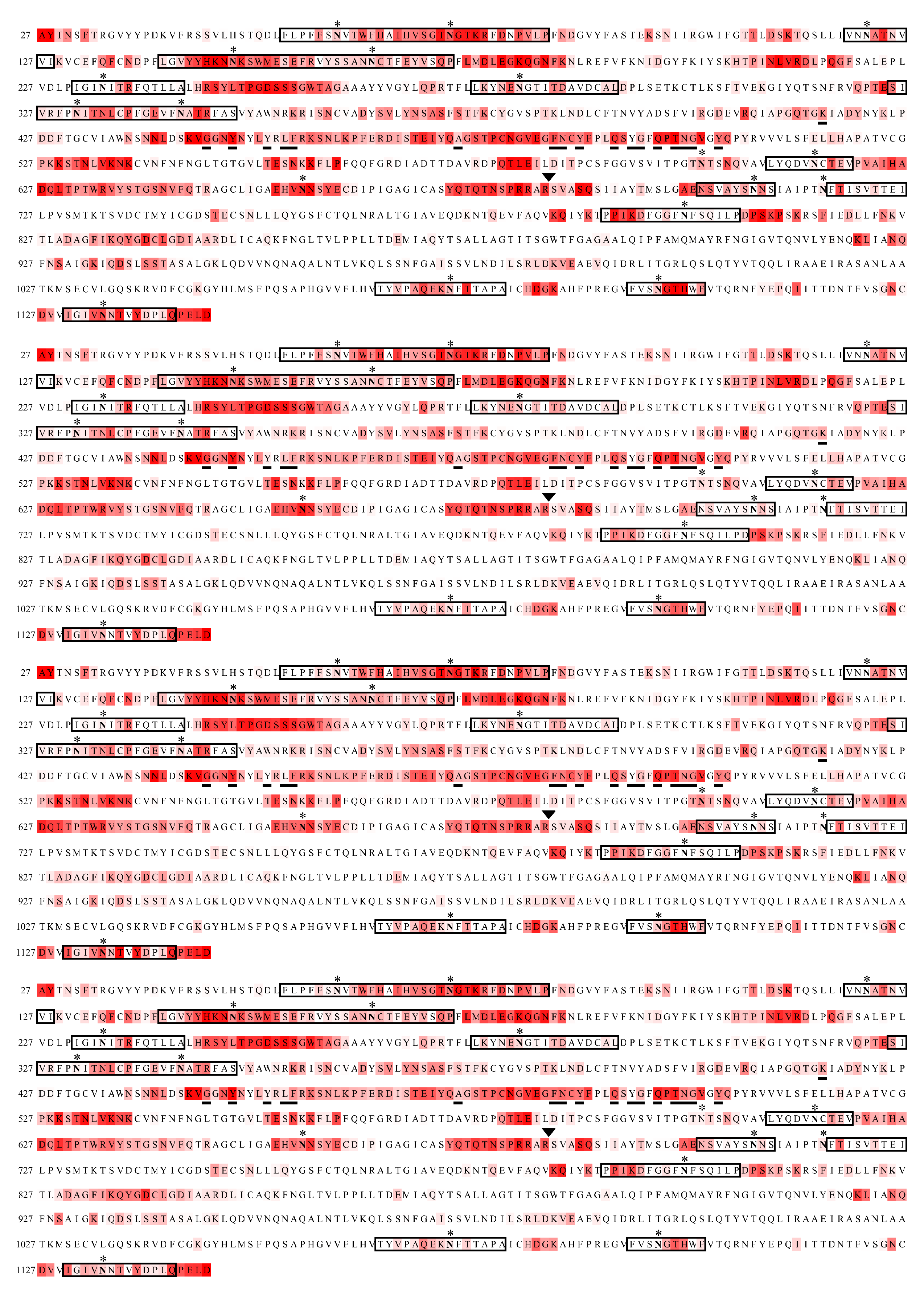

d.

c.

a.

b.

**Supplementary Figure S2.** Sequence of the S protein model colored by antibody accessibility from white to red (0 to 100 % accessible) computed for the M3 (**a.**), M9 (**b**.), Complex **(c.**), and Complex Core F (**d.**) glycoforms. Glycosites are indicated with asterisks, residues reported to interact with the ACE2 receptor are underlined, and the protease cleavage site is indicated with a triangle above the RS junction.

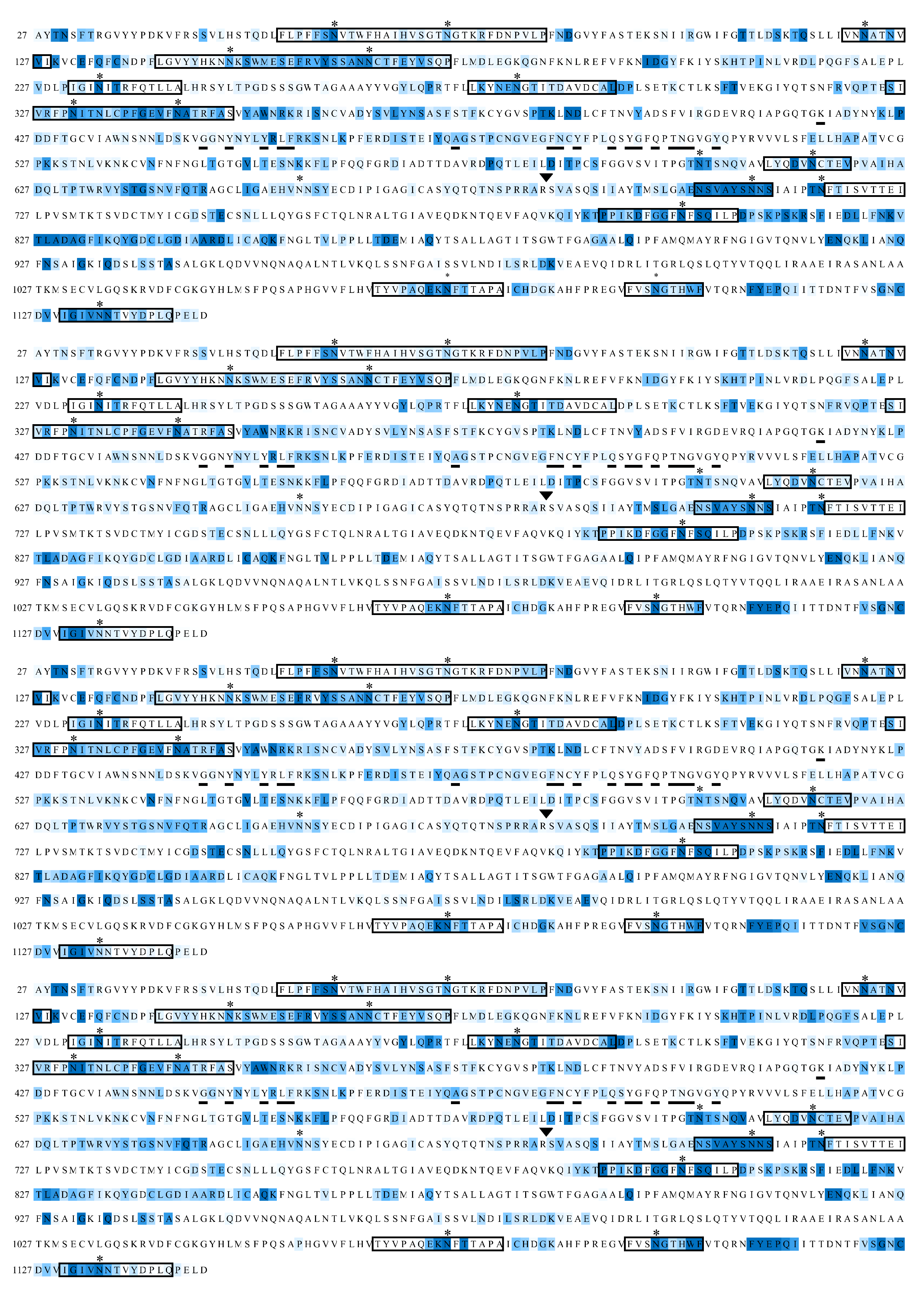

a.

b.

c.

d.

**Supplementary Figure S3.** Sequence of the S protein with the difference in antibody accessibilities between glycoforms plotted as the fold change in accessibility from -4 to 0 (blue to white), where blue indicates glycosylation-dependent surface shielding, computed for the M3 (**a.**), M9 (**b**.), Complex **(c.**), and Complex Core F (**d.**) glycoforms. Glycosites are indicated with asterisks, residues reported to interact with the ACE2 receptor are underlined, and the protease cleavage site is indicated with a triangle above the RS junction.

**Supplementary Table S1.** Structures of the glycans attached at each glycosite in the Site Specific Model

| Hybrid^a^  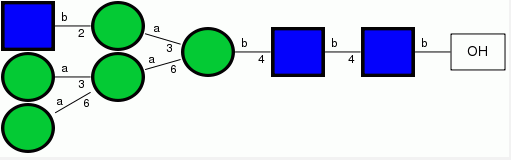 | DManpα1-6[DManpα1-3]DManpα1-6[DGlcpNAcβ1-2DManpα1-3]DManpβ1-4DGlcpNAcβ1-4DGlcpNAcβ1-OH  Glycosite residue: 657 |
| --- | --- |
| FA2  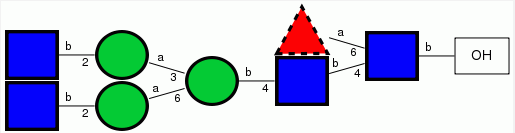 | DGlcpNAcβ1-2DManpα1-6[DGlcpNAcβ1-2DManpα1-3]DManpβ1-4DGlcpNAcβ1-4[LFucpα1-6]DGlcpNAcβ1-OH  Glycosite residues: 149, 165, 331, 343, 616, 1134 |
| A2  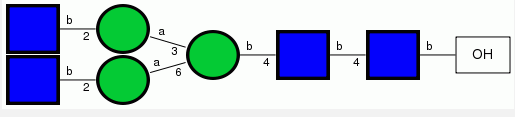 | DGlcpNAcβ1-2DManpα1-6[DGlcpNAcβ1-2DManpα1-3]DManpβ1-4DGlcpNAcβ1-4DGlcpNAcβ1-OH  Glycosite residue: 1098 |
| FA2B  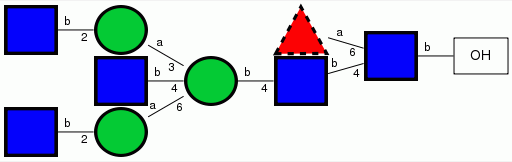 | DGlcpNAcβ1-2DManpα1-6[DGlcpNAcβ1-4][DGlcpNAcβ1-2DManpα1-3]DManpβ1-4DGlcpNAcβ1-4[LFucpα1-6]DGlcpNAcβ1-OH  Glycosite residues: 74, 282 |
| M5  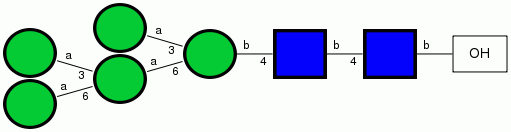 | DManpα1-6[DManpα1-3]DManpα1-6[DManpα1-3]DManpβ1-4DGlcpNAcβ1-4DGlcpNAcβ1-OH  Glycosite residues: 61, 122, 603, 709, 717, 801, 1074 |
| M8  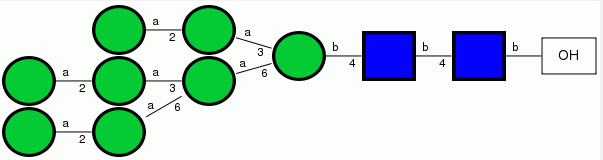 | DManpα1-2DManpα1-6[DManpα1-2DManpα1-3]DManpα1-6[DManpα1-2DManpα1-3]DManpβ1-4DGlcpNAcβ1-4DGlcpNAcβ1-OH  Glycosite residue: 234 |

^a^Glycan naming system from reference 15. In cases where the mass spectrometry data allows for multiple structures, such as FA2/FA1B, the glycan with the least ambiguity was selected. For example, FA2 was selected over FA1B, as the position of the single antennae is ambiguous.

**Supplementary Table S2.** Peptides containing glycosites with sequences predicted^a^ to bind to human HLAs

| HLA Position Specific  Scoring Matrix | RANK | SEQ. POS. | N | SEQUENCE | C | HLA Site | SCORE | % OPT. |
| --- | --- | --- | --- | --- | --- | --- | --- | --- |
| HLA_DR15oDRB1s1501c | 5 | 55 | QDL | FLPFFS**NVT** | WFH | 1 | 15.3 | 36.8 |
| HLA_DR4 | 4 | 58 | FLP | FFS**NVT**WFH | AIH | 1 | 13.3 | 32.2 |
| HLA_DR4oDRB1s0401c | 3 | 58 | FLP | FFS**NVT**WFH | AIH | 1 | 17.6 | 40.0 |
| HLA_DQ7oDQB1s0301c | 13 | 61 | FFS | **NVT**WFHAIH | VSG | 1 | 11.7 | 25.7 |
| HLA_DR8oDRB1s0801c | 6 | 61 | FFS | **NVT**WFHAIH | VSG | 1 | 14.7 | 31.2 |
| HLA_DR1 | 20 | 63 | S**NV** | **T**WFHAIHVS | GTN | 1 | 8.8 | 18.8 |
| HLA_DR4 | 21 | 64 | **NVT** | WFHAIHVSG | TNG | 1 | 10.7 | 25.8 |
| HLA_DR4oDRB1s0401c | 10 | 64 | **NVT** | WFHAIHVSG | TNG | 1 | 12.1 | 27.5 |
| HLA_DR15oDRB1s1501c | 21 | 68 | FHA | IHVSGT**NGT** | KRF | 2 | 10.9 | 26.2 |
| HLA_DR3 | 7 | 75 | GT**N** | **GT**KRFDNPV | LPF | 2 | 11.7 | 29.0 |
| HLA_DR1 | 18 | 76 | T**NG** | **T**KRFDNPVL | PFN | 2 | 9.3 | 19.7 |
| HLA_DR15oDRB1s1501c | 14 | 77 | **NGT** | KRFDNPVLP | FND | 2 | 12.8 | 30.7 |
| HLA_DR15oDRB1s1501c | 7 | 120 | LLI | VN**NAT**NVVI | KVC | 3 | 14.9 | 35.6 |
| HLA_DR8oDRB1s0801c | 2 | 141 | DPF | LGVYYHKN**N** | **KS**W | 4 | 17.1 | 36.2 |
| HLA_DR8oDRB1s0801c | 1 | 142 | PFL | GVYYHKN**NK** | **S**WM | 4 | 20.0 | 42.5 |
| HLA_DR2 | 1 | 143 | FLG | VYYHKN**NKS** | WME | 4 | 22.9 | 45.8 |
| HLA_DR4oDRB1s0402c | 2 | 143 | FLG | VYYHKN**NKS** | WME | 4 | 17.2 | 38.7 |
| HLA_DR5 | 2 | 143 | FLG | VYYHKN**NKS** | WME | 4 | 20.4 | 42.0 |
| HLA_DR1 | 1 | 144 | LGV | YYHKN**NKS**W | MES | 4 | 19.7 | 41.8 |
| HLA_DR11oDRB1s1101c | 19 | 144 | LGV | YYHKN**NKS**W | MES | 4 | 9.7 | 14.7 |
| HLA_DR2 | 8 | 144 | LGV | YYHKN**NK**SW | MES | 4 | 14.4 | 28.8 |
| HLA_DR11oDRB1s1101c | 2 | 145 | GVY | YHKN**NKS**WM | ESE | 4 | 24.7 | 37.6 |
| HLA_DR4 | 18 | 145 | GVY | YHKN**NKS**WM | ESE | 4 | 10.9 | 26.3 |
| HLA_DR4oDRB1s0401c | 4 | 145 | GVY | YHKN**NKS**WM | ESE | 4 | 17.1 | 38.9 |
| HLA_DR7oDRB1s0701c | 2 | 145 | GVY | YHKN**NKS**WM | ESE | 4 | 19.8 | 38.6 |
| HLA_DQ8oDQA1s0301xDQB1s0302c | 2 | 148 | YHK | N**NKS**WMESE | FRV | 4 | 15.2 | 29.7 |
| HLA_DR8oDRB1s0801c | 4 | 157 | ESE | FRVYSSAN**N** | **CT**F | 5 | 15.1 | 32.0 |
| HLA_DR5 | 20 | 159 | EFR | VYSSAN**NCT** | FEY | 5 | 10.5 | 21.7 |
| HLA_DR7 | 1 | 159 | EFR | VYSSAN**NCT** | FEY | 5 | 22.4 | 43.5 |
| HLA_DR1oDRB1s0101c | 8 | 160 | FRV | YSSAN**NCT**F | EYV | 5 | 13.8 | 28.7 |
| HLA_DR3 | 5 | 166 | AN**N** | **CT**FEYVSQP | FLM | 5 | 13.0 | 32.3 |
| HLA_DR4oDRB1s0402c | 13 | 231 | DLP | IGI**NIT**RFQ | TLL | 6 | 11.5 | 25.8 |
| HLA_DR7 | 9 | 234 | IGI | **NIT**RFQTLL | ALH | 6 | 13.0 | 25.3 |
| HLA_DR1oDRB1s0101c | 20 | 235 | GI**N** | **IT**RFQTLLA | LHR | 6 | 9.9 | 20.6 |
| HLA_DR4oDRB1s0401c | 7 | 235 | GI**N** | **IT**RFQTLLA | LHR | 6 | 14.5 | 32.9 |
| HLA_DR4oDRB1s0402c | 11 | 235 | GI**N** | **IT**RFQTLLA | LHR | 6 | 11.7 | 26.3 |
| HLA_DR4 | 12 | 277 | TFL | LKYNE**NGT**I | TDA | 7 | 11.9 | 28.8 |
| HLA_DR5 | 15 | 278 | FLL | KYNE**NGT**IT | DAV | 7 | 11.3 | 23.3 |
| HLA_DR1oDRB1s0101c | 22 | 285 | **NGT** | ITDAVDCAL | DPL | 7 | 9.6 | 20.0 |
| HLA_DR8oDRB1s0801c | 11 | 325 | PTE | SIVRFP**NIT** | NLC | 8 | 13.1 | 27.9 |
| HLA_DR15oDRB1s1501c | 3 | 326 | TES | IVRFP**NIT**N | LCP | 8 | 15.9 | 38.1 |
| HLA_DR4 | 1 | 326 | TES | IVRFP**NIT**N | LCP | 8 | 16.2 | 39.2 |
| HLA_DR15oDRB1s1501c | 22 | 335 | ITN | LCPFGEVF**N** | **AT**R | 9 | 10.6 | 25.4 |
| HLA_DR15oDRB1s1501c | 11 | 341 | FGE | VF**NAT**RFAS | VYA | 9 | 13.4 | 32.0 |
| HLA_DQ7oDQB1s0301c | 10 | 611 | VAV | LYQDV**NCT**E | VPV | 10 | 13.1 | 28.8 |
| HLA_DR7oDRB1s0701c | 3 | 612 | AVL | YQDV**NCT**EV | PVA | 10 | 16.2 | 31.6 |
| HLA_DR8oDRB1s0801c | 14 | 703 | GAE | NSVAYS**NNS** | IAI | 11 | 12.5 | 26.5 |
| HLA_DR4oDRB1s0401c | 18 | 718 | PT**N** | **FT**ISVTTEI | LPV | 12 | 10.1 | 22.9 |
| HLA_DR5 | 12 | 792 | YKT | PPIKDFGGF | **NFS** | 13 | 11.7 | 24.1 |
| HLA_DQ8oDQA1s0301xDQB1s0302c | 3 | 794 | TPP | IKDFGGF**NF** | **S**QI | 13 | 15.0 | 29.3 |
| HLA_DR4 | 11 | 797 | IKD | FGGF**NFS**QI | LPD | 13 | 12.1 | 29.3 |
| HLA_DR15oDRB1s1501c | 12 | 799 | DFG | GF**NFS**QILP | DPS | 13 | 13.1 | 31.4 |
| HLA_DR2 | 5 | 1066 | LHV | TYVPAQEK**N** | **FT**T | 14 | 15.3 | 30.6 |
| HLA_DR7 | 5 | 1066 | LHV | TYVPAQEK**N** | **FT**T | 14 | 17.2 | 33.4 |
| HLA_DR4 | 7 | 1072 | PAQ | EK**NFT**TAPA | ICH | 14 | 12.9 | 31.2 |
| HLA_DR4 | 2 | 1095 | EGV | FVS**NGT**HWF | VTQ | 15 | 13.4 | 32.4 |
| HLA_DR4oDRB1s0401c | 2 | 1095 | EGV | FVS**NGT**HWF | VTQ | 15 | 17.7 | 40.2 |
| HLA_DR7oDRB1s0701c | 5 | 1095 | EGV | FVS**NGT**HWF | VTQ | 15 | 15.0 | 29.2 |
| HLA_DR2 | 20 | 1130 | DVV | IGIV**NNT**VY | DPL | 16 | 12.5 | 25.0 |
| HLA_DR8oDRB1s0801c | 15 | 1135 | IV**N** | **NT**VYDPLQP | ELD | 16 | 12.3 | 26.0 |
| HLA_DR2 | 17 | 1154 | ELD | KYFK**NHT**SP | DVD | 17 | 12.7 | 25.4 |
| HLA_DR5 | 5 | 1154 | ELD | KYFK**NHT**SP | DVD | 17 | 15.9 | 32.8 |
| HLA_DR7 | 6 | 1154 | ELD | KYFK**NHT**SP | DVD | 17 | 16.8 | 32.7 |
| HLA_DR4oDRB1s0401c | 1 | 1155 | LDK | YFK**NHT**SPD | VDL | 17 | 20.1 | 45.6 |
| HLA_DR51oDRB5s0101c | 4 | 1155 | LDK | YFK**NHT**SPD | VDL | 17 | 16.8 | 42.6 |
| HLA_DR7oDRB1s0701c | 12 | 1155 | LDK | YFK**NHT**SPD | VDL | 17 | 12.1 | 23.5 |
| HLA_DR4oDRB1s0405c | 4 | 1176 | **NAS** | VVNIQKEID | RLN | 18 | 15.0 | 35.8 |
| HLA_DQ8oDQA1s0301xDQB1s0302c | 7 | 1187 | DRL | NEVAKNL**NE** | **S**LI | 19 | 12.8 | 25.0 |
| HLA_DR8oDRB1s0801c | 16 | 1187 | DRL | NEVAKNL**NE** | **S**LI | 19 | 11.1 | 23.5 |

^a^Output from the Rankpep server (imed.med.ucm.es/Tools/rankpep)
